# Quantifying the impact of rare and ultra-rare coding variation across the phenotypic spectrum

**DOI:** 10.1101/148247

**Authors:** Andrea Ganna, Kyle F. Satterstrom, Seyedeh M Zekavat, Indraniel Das, Mitja I. Kurki, Claire Churchhouse, Jessica Alfoldi, Alicia R. Martin, Aki S. Havulinna, Andrea Byrnes, Wesley K. Thompson, Philip R. Nielsen, Konrad J. Karczewski, Elmo Saarentaus, Manuel A. Rivas, Namrata Gupta, Olli Pietiläinen, Connor A Emdin, Francesco Lescai, Jonas Bybjerg-Grauholm, Jason Flannick, on behalf of GoT2D/T2D-GENES consortium, Josep Mercader, Miriam Udler, on behalf of SIGMA consortium, Helmsley IBD Exome Sequencing Project, FinMetSeq Consortium, iPSYCH-Broad Consortium, Markku Laakso, Veikko Salomaa, Christina Hultman, Samuli Ripatti, Eija Hämäläinen, Jukka S Moilanen, Jarmo Körkkö, Outi Kuismin, Merete Nordentoft, David M. Hougaard, Ole Mors, Thomas Werge, Preben Bo Mortensen, Daniel MacArthur, Mark J. Daly, Patrick F. Sullivan, Adam E. Locke, Aarno Palotie, Anders D. Børglum, Sekar Kathiresan, Benjamin M. Neale

## Abstract

There is a limited understanding about the impact of rare protein truncating variants across multiple phenotypes. We explore the impact of this class of variants on 13 quantitative traits and 10 diseases using whole-exome sequencing data from 100,296 individuals. Protein truncating variants in genes intolerant to this class of mutations increased risk of autism, schizophrenia, bipolar disorder, intellectual disability, ADHD. In individuals without these disorders, there was an association with shorter height, lower education, increased hospitalization and reduced age. Gene sets implicated from GWAS did not show a significant protein truncating variants-burden beyond what captured by established Mendelian genes. In conclusion, we provide the most thorough investigation to date of the impact of rare deleterious coding variants on complex traits, suggesting widespread pleiotropic risk.

**Main abbreviations:**

PTV: = Protein Truncating Variants
PI: = Protein Truncating Intolerant
PI-PTV: = Protein Truncating Variant in genes that are Intolerant to Protein Truncating Variants

## Main text

Protein truncating variants (PTVs) are likely to modify gene function and have been linked to hundreds of Mendelian disorders ^1,2^. However, the impact of PTVs on complex traits has been limited by the available sample size of whole-exome sequencing studies (WES) ^3^. Here, we assembled whole-exome sequencing data from 100,296 individuals, drawing from a combination of cohort and case/control disease studies with phenotypic information on a total of 13 quantitative traits and 10 diseases (**Supplementary Table 1-3**). We used a common pipeline to process, annotate and analyze the data (**Supplementary Material** and **Supplementary Figure 1** for principal components plots).

We began by focusing our analysis on PTVs that occur in a set of 3,172 PTV-intolerant (PI) genes (**Supplementary Table 4** reports all gene sets used in this study). Our motivation for focusing on the PI-PTVs was two-fold. First, this gene class was identified through an unbiased approach that leveraged the observed frequency distribution in ExAC ^4^ without relying on information from model organisms or *in vitro* experiments. Second, PI-PTVs have been shown to associate with early onset neurodevelopmental and psychiatric disorders, and are likely to result in reproductively disadvantageous phenotypes ^5 6,7^. To focus on those variants that are most likely to be subject to purifying selection, we considered only rare (allele frequency < 0.1%) and ultra-rare (observed in less than 1 in 201,176 individuals) variants (**Supplementary Materials**).

After excluding participants diagnosed with a psychiatric or neurodevelopmental disorder, we observed an average of 7.72 and 0.30 rare PTVs per individual, across all genes and in PI genes respectively (Figure 1a); one or more ultra-rare PI-PTV was observed in 11% of the individuals. The number and frequency of rare variants differs across populations, reflecting the degree of selection compounded by recent demography, including bottlenecks, split times, and migration between populations ^8^. The ratio of deleterious to neutral alleles per individual increases as humans migrated out of Africa, consistent with less efficient negative selection against deleterious variants and serial founder effects that reduce the effective population size ^9^. Conditional on a variant being ultra-rare, we observe a higher ratio of PTV to synonymous variants (Figure 1b); recently arisen ultra-rare variants have had less time to be purged by negative selection, which is further magnified in populations that have undergone a recent bottleneck. For example, we observed a higher ratio among Ashkenazi Jewish and Finnish populations as compared to non-Finnish Europeans, reflecting the more recent population-specific bottlenecks ^10,11^.

**Fig. 1.**
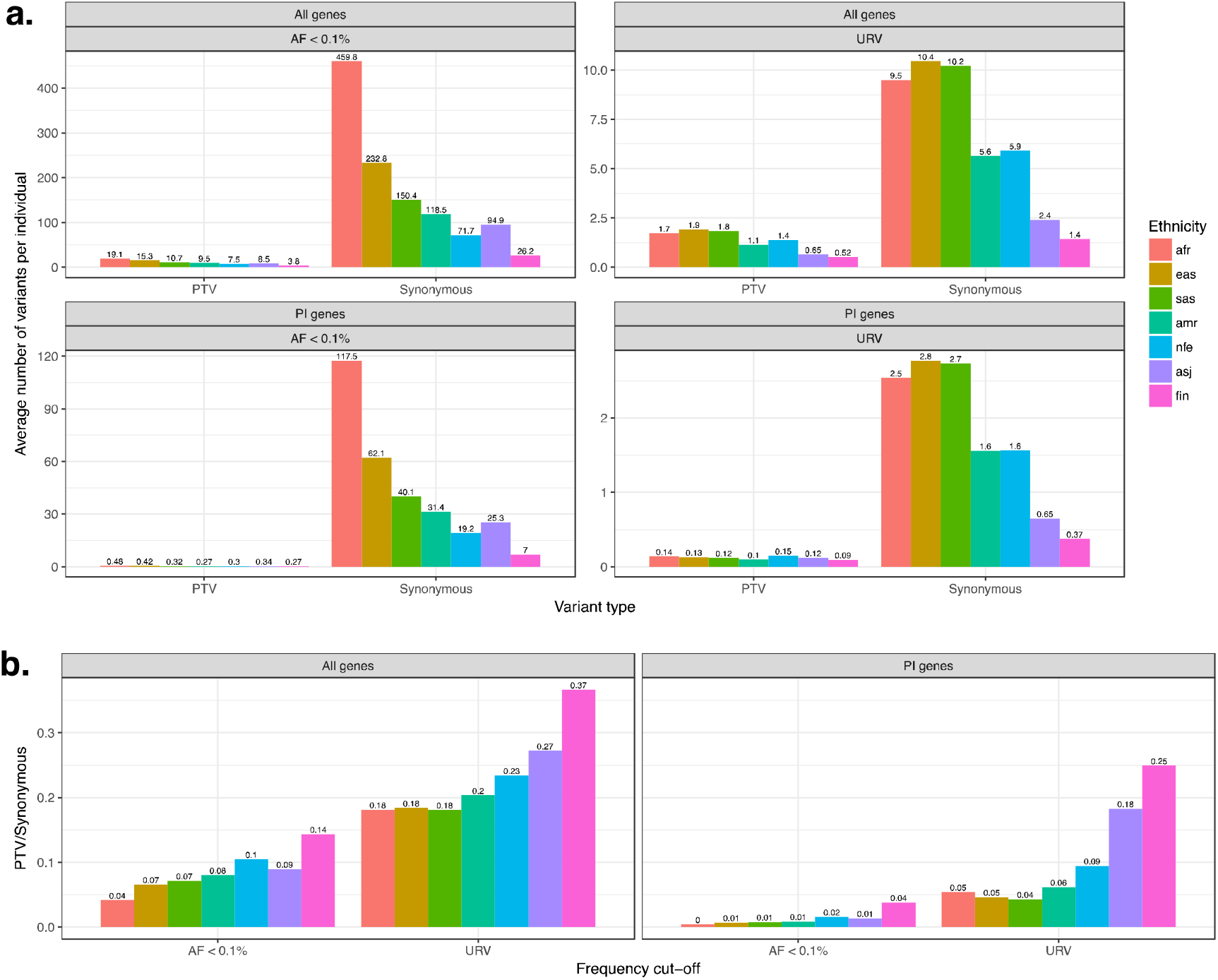
**A.** Average number of variants per individual in N=83,439 participants without neurodevelopmental/psychiatric disorders. We report the results separately for each ethnic group. **B.** Ratio between PTV/Synonymous for each ethnic group. *Afr*: African American, *Eas*: East Asian, *Sas*: South Asian, *Amr*: Latinos, *Nfe*: Non Finnish European, *Asj*: Ashkenazi Jewish, *Fin*: Finnish.

We tested the association between a burden of PI-PTVs and the 13 traits and 10 disease diagnoses (Figure 2) by performing study and ethnicity-specific linear or logistic regression analysis adjusting for potential confounders such as overall mutation rate (**Supplementary Table 5**). The results of these separate analyses were then meta-analyzed (**Supplementary Materials**). We used an experiment-wise p-value threshold of 2x10^−3^ to account for multiple testing (0.05/23 traits tested). Among the quantitative traits, we found that carriers of at least one rare PI-PTV had fewer years of education (-2.2 months, p=4x10^−4^), as we have previously reported ^12^, were shorter (-0.2 cm, p=3x10^−4^), and younger (-3.7 months, p=2x10^−7^).

**Fig. 2.**
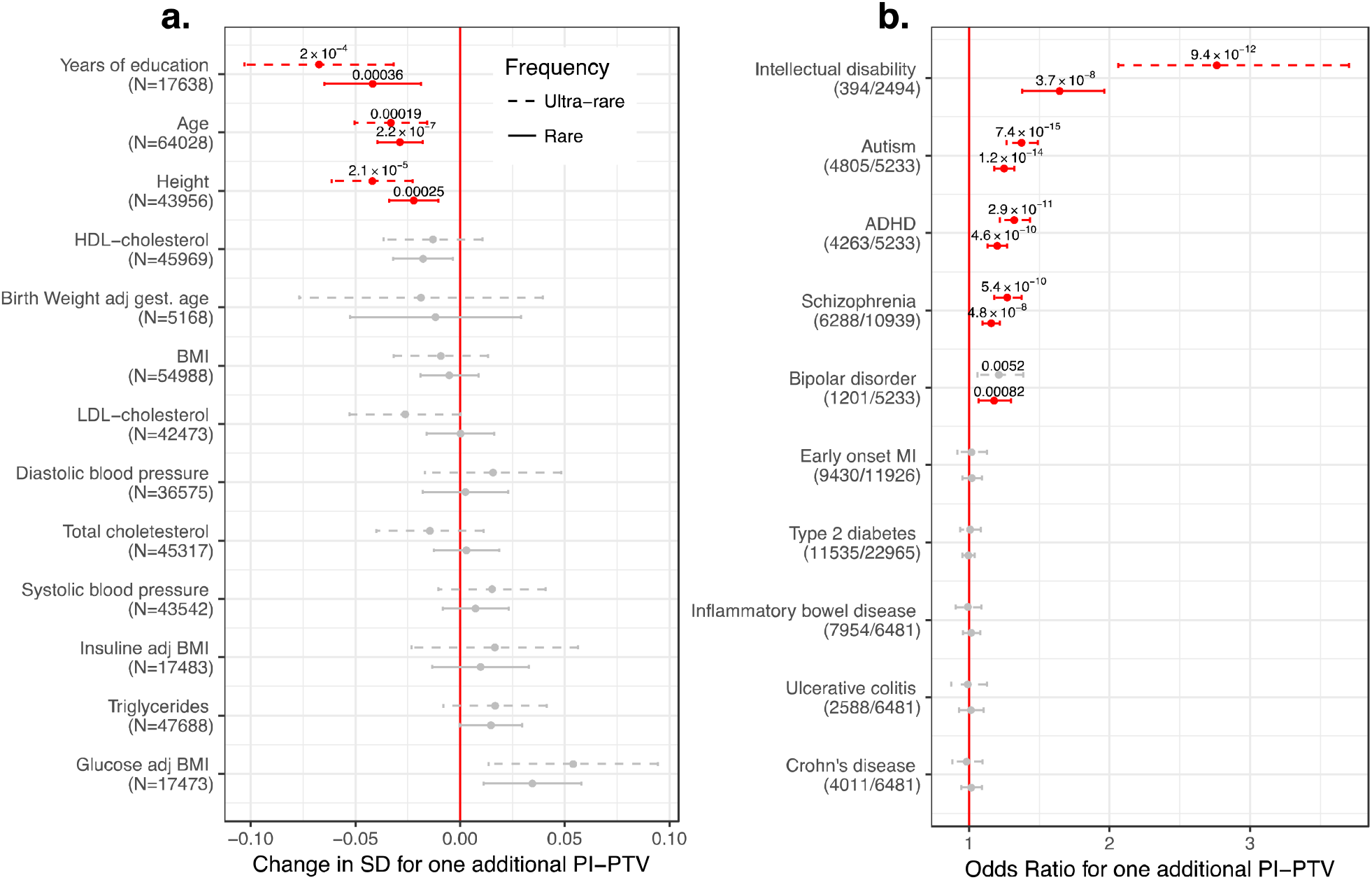
**A.** Association between PI-PTV burden and continuous traits. We reported the association in standard deviations (SD) to allow for comparison across traits. In brackets, we reported the number of individual included in the analysis for each trait. The P-values are reported only for experiment-wise significant results (p < 2x10^−3^), highlighted in red. Bars indicate 95% confidence intervals. All the results are obtained from meta-analyzing study and ethnicity-specific associations. **B.** Odds ratio for association between PI-PTV burden and dichotomous traits. In brackets, we reported the number of cases and controls.

To ensure the robustness of the age result, we performed a series of quality control analyses to guard against the impact of technical confounders or specific study designs that might bias the results. We first confirm that the signal was not observed among PTVs in non-PI genes and synonymous variants in PI genes, our negative controls (**Supplementary Figure 2**). We further found that the effect was consistent across ethnicities and study cohorts (**Supplementary Figure 3**), when INDELs and SNPs were considered separately, when mutations possibly caused by cytosine deamination (C ->T or G -> A) were excluded and when a highly stringed QC was used (**Supplementary Figure 4**). Similarly, the association did not change after adjusting for 8 QC metrics capturing most of the sample properties (**Supplementary Figure 5**). Although we could not exclude the impact of unmeasured confounder, we find this result consistent with reduced survival, detrimental health or decreased study participation over time among PI-PTVs carries. If reduced survival or detrimental health effects drive this association, we see it as a signature of viability selection, overall in the population.

**Fig. 3.**
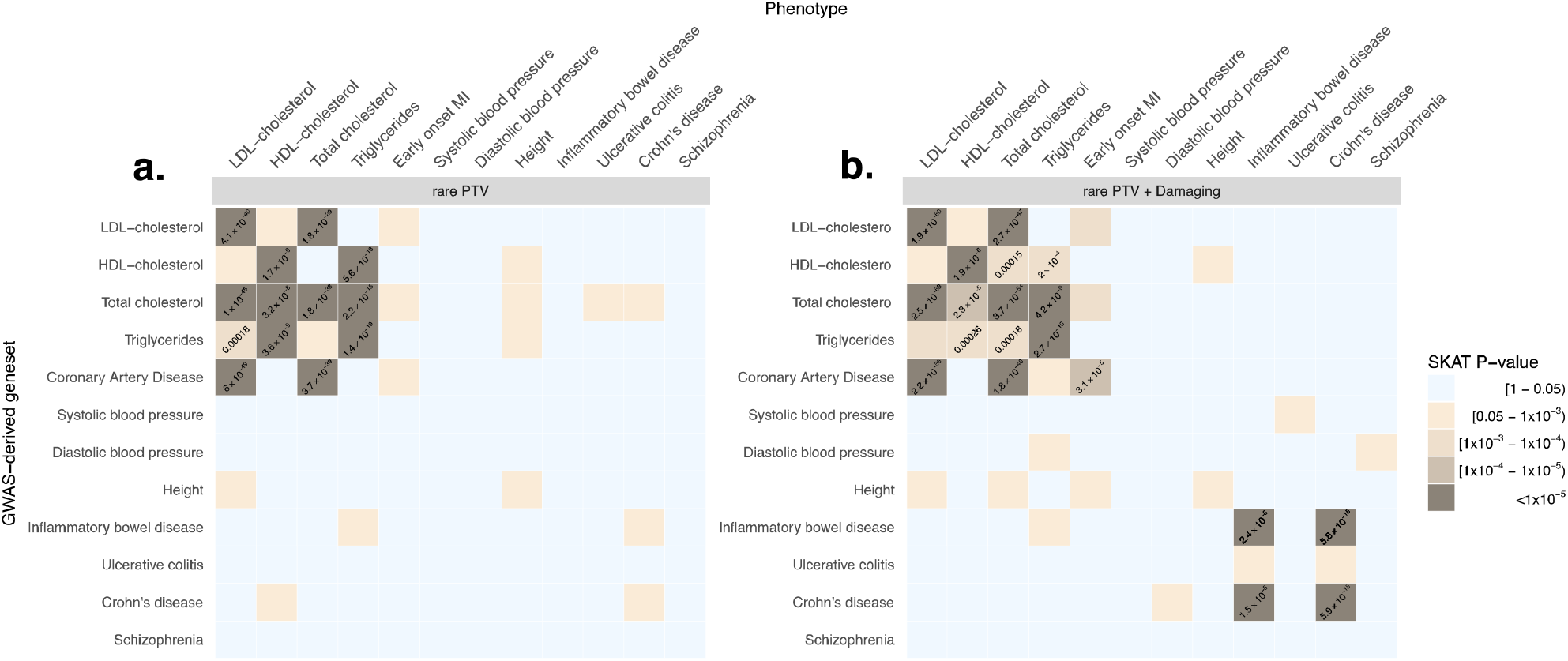
**A**. Association (SKAT test P-value) in GWAS-derived gene sets (y axis) between rare PTVs and the phenotypes reported on the x axis. Each geneset is obtained using DEPICT to link SNPs derived from GWAS with P-value < 5x10^−8^ and a candidate gene. In brackets we report the number of genes with at least one PTV in our dataset. P-values are reported only for experiment-wise associations (p < 0.0003). **B.** Association (SKAT test P-value) in GWAS-derived gene sets (y axis) between rare PTVs + damaging missense and the phenotypes reported on the x axis.

We then focused on dichotomous traits (Figure 2). We observed significant associations with all psychiatric disorders that were tested – intellectual disability (ID) (odds ratio [OR]=1.7, p=4x10^−8^), autism (OR=1.3,p=1x10^−14^), schizophrenia (OR=1.2,p=5x10^−8^), ADHD (OR=1.2, p=5x10^−10^) and bipolar disorder (OR=1.2, p=8x10^−4^). We did not however find PI-PTV burden to be associated with later onset, non-brain-related diseases such as type 2 diabetes, early onset myocardial infarction, inflammatory bowel disease, ulcerative colitis or Crohn’s disease. Across all significantly associated phenotypes, the effect size was stronger among the subset of ultra-rare PI-PTV carriers, confirming that rarer PTVs are, on average, more deleterious.

The association with these five neurodevelopmental/psychiatric disorders and three quantitative traits was only observed for PI-PTVs and not for PTVs in non-PI genes nor for synonymous variants in PI genes. These results suggest that the association to PI-PTVs is not driven by population stratification or technical bias (**Supplementary Figures 6**).

Our approach so far focuses on assuming that all PI-PTVs act on the phenotype in the same direction, that is, they are all either protective or risk conferring. We relaxed this hypothesis, allowing rare PI-PTVs to have different directions as well as different magnitudes of effects, and repeated these tests using SKAT ^13^. We did not identify any additional associations (**Supplementary Figure 7**), suggesting that PI genes do not account for a substantial fraction of variability in the traits for which no PTV burden was identified. Further, the observed burden of PI-PTVs for neurodevelopmental/psychiatric disorders, height, educational attainment and age suggests that the majority of those PI genes that have an effect, do so in the same direction. Although case ascertainment bias reduces the power of detecting protective variants, this is not the case for continuous traits like height.

We also evaluated whether damaging missense variants, which are on average more common and less severely deleterious than PTVs, showed a similar signal. Damaging missense variants have been associated with complex disorders such as coronary heart disease and inflammatory bowel disease^14 15^. We found an independent signal for damaging missense variants in PI genes for all disorders and traits that were also associated with PI-PTVs. Furthermore, the strength of the association increased as a function of the number of prediction algorithms that confidently classified a missense variant as ‘damaging’ (**Supplementary Figure 8**) suggesting that these missense mutations are similar to PTVs in biological effect, potentially abrogating gene function. We note that this effect was particularly strong for ultra-rare variants, reinforcing the observation that variant frequency is a marker of selection and aids in the identification of pathogenic damaging missense variation^16 17^.

Given the high degree of shared comorbidities across neurodevelopmental/psychiatric disorders, we leveraged information from the Danish National Psychiatric registry to evaluate whether the signal was driven by a specific disorder or shared across multiple disorders. Individuals with multiple neurodevelopmental/psychiatric disorders, and especially those with ID, showed a stronger enrichment of PI-PTVs (**Supplementary Figure 9**). Nevertheless, among those without comorbidities, the signal remained significant and remarkably similar across disorders (OR=1.12, 1.15, 1.21, 1.18 for schizophrenia, bipolar, autism and ADHD, respectively; Cochran’s Q test for heterogeneity P = 0.282). We further found that carriers of ultra-rare PI-PTVs had earlier onset of ADHD (-4.0 months, p =0.008; **Supplementary Table 6**). However, this was partially explained by the fact that individuals with earlier diagnosis of ADHD were also more likely to be diagnosed with ID (14.7 vs. 15.5 years for ADHD patients with and without ID, T-test P-value=0.009). Indeed when we considered ADHD cases without major comorbidities, the effect was attenuated (-2.9 months, p=0.12). Finally, in controls with none of these psychiatric diagnoses, we still observed a significant association with the broader ICD-10 category of mental, behavioral and neurodevelopmental disorders, suggesting that PI-PTVs influence the broader cognitive spectrum (**Supplementary Table 7**).

Since previous studies have shown a higher rate of PI *de novo* PTVs in autism-affected females as compared to males ^18 19^, we wondered if sex played a role here. In this study however, we did not have parent-offspring subjects needed to distinguish *de novo* variants from those that have recently arisen in the population, the latter being the majority of observed rare variants. This would potentially dilute the sex-specific effect if it is in fact a property of *de novo* variants but not of rare variants more generally. We found both weak and insignificant differences between males and females in the effect of PI-PTVs on 4 neurodevelopmental/psychiatric disorders (**Supplementary Table 8**). Interestingly, we did not observe differences in ADHD-affected males and females, in contrast with the hypothesis that affected females might be enriched for rare deleterious variants^20^. We cannot exclude that differences in the diagnostic criteria used in these European studies compared to those of previous studies, which were mostly conducted in the U.S., might explain these results.

We also assessed whether the observed burden of PTVs was specific to PI genes, or such a burden could be identified for other gene sets that are likely to contain functionally relevant genes. First, we examined other experimental and literature-based gene sets linked to severe phenotypes. Specifically, we considered all genes (i) reported in ClinVar ^2^, (ii) that resulted in lethal or subviable phenotypes in mice ^21^, (iii) that were required for proliferation and survival in a human cancer cell line ^22^ and those (iv) that were categorized as haploinsufficient by ClinGen (**Supplementary Table 4** and **Supplementary Materials**). Except for the haploinsufficient genes, which showed a significantly stronger association with autism alone, none of the other gene sets tested showed the PTV burden that was captured by PI genes (**Supplementary Figure 10**). This suggests that the degree of natural selection against PTVs in a gene is indeed an important indicator of whether such PTVs are likely to be implicated as strong effects for neurodevelopmental/psychiatric disorders, height, educational attainment and age. It also highlights that observed associations are not simply reflecting an aggregate signal from known Mendelian disorders. Indeed we could not detect significant association for any of the traits considered in this analysis when just focusing on ClinVar genes.

We reasoned that a single variant approach, rather than a gene-based test, might provide increased resolution. We considered all high quality ClinVar variants (0.76 on average per individual) and a set of variants deemed to be recessive lethal (0.03) (**Supplementary Materials**). Carriers of these variants were not enriched in any of the disorders or traits examined here (**Supplementary Table 9**).

Second, we examined whether results from GWAS conducted on the same phenotypes as those included in this study could implicate genes containing an aggregate PTV burden. We used DEPICT^23^ to link genome-wide significant hits to candidate genes (**Supplementary Table 10** and **Supplementary Materials**) and, within each GWAS-derived gene set, we studied the association between rare PTVs and the phenotypes using the SKAT test. GWAS-derived gene-sets captured associations between rare PTVs and different classes of lipids (Figure 3 and **Supplementary Table 11**). For example, the association between rare PTVs and HDL-cholesterol was captured by gene sets derived from GWAS of HDL (p=2x10^−9^), total cholesterol (p=3x10^−8^) and triglycerides (p=4x10^−9^), but not by those of coronary heart diseases (p=0.25), consistent with previous observations about non-causality of HDL-cholesterol on coronary heart diseases ^24^. The inclusion of both rare damaging missense and PTVs resulted in additional signal co-localization between inflammatory bowel disease, early onset myocardial infarction and the corresponding GWAS-derived gene-sets. However, it appeared that all these signals were being driven by well-known genes, involved in rare familial forms of these diseases. Specifically, when Mendelian lipid genes and *NOD2* were removed from the cardiovascular and inflammatory bowel disease-related GWAS gene-sets respectively, no signal remained (**Supplementary Figure 11**). This might reflect a lack of power (despite this being the largest WES study for the majority of the traits), inaccurate links between genome-wide significant hits and the corresponding candidate genes or PTVs and common variants acting on partially distinct pathways. Nevertheless, we observed similar results when including SNPs below genome-wide significance to increase power and when using different methods to link SNPs with corresponding candidate genes to increase precision, including gene-based testing ^25^ and eQTL mapping **(Supplementary Figures 11** and **Supplementary Materials)**.

The choice of the 10 diseases and 13 quantitative traits included in the main analysis was driven by data availability and power considerations, but was not truly unbiased. Therefore, we leveraged national population health registries to increase the scope of disorders we could examine. These well-studied and validated registries ^26,27^ include diagnostic codes from 14,117 individuals (N=8,493 from Finland and 5,624 from Sweden), recorded between 1968 to 2015 (**Supplementary Table 12**). Individuals with psychiatric disorders were excluded from our analyses. To maximize the validity of the diagnoses, we used a curated list of disease definitions aggregating related ICD codes (**Supplementary Table 13**). We studied the association between rare PI-PTVs and 101 diseases with at least 50 cases, using a survival analysis model. We identified a novel association (multi-testing significance threshold = 0.05/101; 5x10^−4^) with chronic kidney failure (hazard ratio=1.9, p=3x10^−6^; number of cases=120; **Supplementary Figure 12**). The association was strong among the Finnish data and only significant when considering ultra-rare PI-PTVs in the Swedish data (**Supplementary Table 14**).

We speculated that this association might reflect a burden of underlying comorbidities that were too rare to be included in this analysis. To evaluate epidemiological associations, we extended our analysis to 28,709 Finnish individuals that were not exome-sequenced, but were linked to the registries. We found that patients with chronic kidney failure also have a higher rate of cardiovascular-related comorbidities, as well as skin infections, kidney cancer and other abnormalities of the renal system (**Supplementary Table 15**). Therefore, it is challenging, to determine whether it is the chronic kidney failure or some more rare comorbid condition that drives the association with PI-PTVs.

We also examined whether the association between PI-PTVs and diminished cognition and detrimental health would result in a higher number of hospital visits, counting the number of inpatient visits associated with a unique ICD codes. In both the Swedish and Finnish datasets, we observed a significant increase in the rate of hospital visits with a greater burden of PI-PTVs (+7.6% per additional PI-PTV, p=0.0002). We used different strategies to model the outcome and observed similar results (**Supplementary Table 16** and **Supplementary Figure 13**).

By aggregating WES data on more than 100,000 individuals for 23 different traits and disorders, we have gained insight into the role of PTVs in conferring risk for these conditions. First, PTVs occurring in PI genes had a remarkably similar effect on autism, schizophrenia, bipolar disorder and, for the first time, we showed a similar effect on ADHD. The observed effects were not driven by major underlying comorbidities. This suggests that these PI-PTVs as a whole are likely to be pleiotropic, influencing some core intermediate phenotypes that relate to risk across many psychiatric disorders. Nevertheless, we could only consider `bulk` pleiotropy, which is the combined impact of PTVs in PI genes and we are not powered to detect if single variants have disease-specific effects. Further, this burden suggests that individual PI genes will be eventually discovered conclusively for each of these disorders, not just autism, but such associations will need to be interpreted in the light of this shared effect across disorders. The strong enrichment of PI-PTVs in individuals with neurodevelopmental/psychiatric disorders does not exclude the existence of non-PI genes involved in the etiology of these disorders. These genes, however are more likely to have weaker and, possibly, trait-specific effect.

Second, we detected for the first time a significant association between PI-PTVs and decreased human height. In contrast to this, a recent large-scale study using the exome-chip has shown a similar numbers of height-increasing and height-decreasing rare variants ^28^. This discrepancy could be because, by using a more stringent frequency cut-off and focusing on a subset of genes likely to cause early onset severe disease, we effectively considered variants related to a burden of (incompletely) penetrant Mendelian-type disorders, often characterized by reduced growth. Such an interpretation is consistent with a tighter link to directional selection on stronger impact mutations for human height.

Third, we systematically compared the co-localization of signal between GWAS-candidate genes and rare PTVs. We found few overlaps (cardiovascular-related traits, inflammatory bowel disease) which, we revealed, were entirely driven by a few genes previously identified by both GWAS and WES studies. Other traits did not show any overlap. Schizophrenia, for example, which is highly enriched for PI-PTVs, did not show overlap with GWAS candidate genes. Even among traits where genes with low-frequency coding variants have been previously identified by exome-chip-based studies, such as height and systolic blood pressure, we found no substantial rare PTVs enrichment. These results suggest that the relationship between GWAS signal and rare coding variants is not always straightforward, and that, when interpreting WES data, other complementary approaches such as those that integrate population genetic models and large sample resources, might be more suitable to nominate gene sets of interest. The degree of overlap, and therefore the most effective strategy to identify pathogenic variants, is likely to depend on the selective pressure shaping the genetic architecture of the trait under investigation. Moreover, it cannot be overlooked that individuals carrying rare PTVs in genes implicated by common variant-based approaches might present phenotypic outcomes that deviate from those under investigation. Finally, it is interesting to notice that, while PTVs tend to have a consistent directional effect within PI gene, this is not the case for GWAS-derived gene sets, where most of signals could only be captured by assuming heterogeneity in effect direction (**Supplementary Notes**).

In conclusion, in the largest WES study to date, we showed that PI genes are well suited to capture the impact of rare to ultra-rare PTVs on the cognitive, behavioral and developmental spectra. This is less the case for major later-onset complex traits with modest effect on reproductive fitness. Strategies to prioritize gene sets relevant for these traits would need to consider the role that relaxed selective pressure has been playing in shaping the frequency distribution of disease-causing PTVs.

## Acknowledgments

A.G. is supported by the Knut and Alice Wallenberg Foundation (2015.0327) and the Swedish Research Council (2016-00250). This study was supported by grants from the National Human Genome Research Institute (U54 HG003067, R01 HG006855), the National Institute of Mental Health (1U01MH105666-01, 1R01MH101244-02), the National Institute of Diabetes and Digestive and Kidney Disease (1U54DK105566-02), the Stanley Center for Psychiatric Research, the Alexander and Margaret Stewart Trust, the National Institutes of Mental Health (R01 MH077139 and RC2 MH089905) and the Sylvan C. Herman Foundation. V.S. was supported by the Finnish Foundation for Cardiovascular Research.

## Code and data sharing

We have made the code and variants used in the paper available at: https://github.com/andgan/ultra_rare_pheno_spectrum

We are extremely committed to data sharing. However, we recognize that due to national regulation not all individual level data and phenotypes can be shared using dbGap. Specifically, access to the Danish and Finnish phenotypic and genetic data can only be obtained by Danish and Finnish national institutions. Information about getting access to the Danish data can be obtained at http://ipsych.au.dk/about-ipsych/. Access to the Finnish data can be obtained at https://www.thl.fi/en/web/thl-biobank/for-researchers. We are happy to help other researchers with information to simplify the application process to obtain these datasets. All the remaining datasets are available via dbGap. We are currently trying to release the entire dataset in dbGap, but if that would not be possible, we will provide a detailed description on the procedures to access the individual datasets that contributed to this study.

